# Dog population estimates and rabies vaccination optimization using the High-Resolution Settlement Layer (HRSL) – a proof of principle for the Oshana region, Namibia

**DOI:** 10.1101/2023.05.11.540335

**Authors:** Conrad M. Freuling, Eva-Maria Czaja, Patrick Wysocki, Rauna Athingo, Tenzin Tenzin, Thomas Müller

## Abstract

Despite being vaccine preventable, dog-mediated rabies continues unabated in low-resourced countries in Africa and Asia. For interventions into dog rabies control, an estimate of the dog population is a prerequisite. Here we used a High-Resolution Settlement Layer (HRSL) with an unprecedented resolution of 30m grid length that is Open Source for dog populations estimates and studies on vaccination coverages, with the Oshana region of Namibia as an example. Our analyses show that the average dog density per km^2^ is 8.15 but ranges between 0 and 40 per constituency, with individual densities being as high as 551.

Spatial analyses for different settings of static vaccination points indicate that the previously used vaccination points during the pilot phase and cattle crush-pens are insufficient for reaching a 70% vaccination level in the Oshana region. Based on cost calculations, between US$5.29 and US$7.77 are needed to parenterally vaccinate dogs in this region, suggesting that oral rabies vaccination may be a cost-effective supplement or even replacement. The high-resolution spatial analyses are exemplified for rabies, but any other One Health intervention, particularly for Neglected tropical diseases in highly heterogenous and remote areas could use our approach as a template.

**Author Summary:** Here, we used high-resolution geospatial data for demonstrating and validating its utility to assist veterinary authorities in their fight against dog-mediated rabies on the example of the Oshana region in Namibia. With such detailed data it is possible estimate the dog population and to analyse and optimize vaccination strategies in dog rabies endemic areas. Such analyses are exemplified for rabies, but any other One Health intervention, particularly for Neglected tropical diseases in highly heterogenous and remote areas could use our approach as a template.

## 1. Introduction

Rabies, a viral zoonotic disease, is still causing an estimated 59,000 human deaths annually (95% CI 25,000–159,000) [1], despite the availability of biologicals for humans and vaccines for dogs. Dogs are the main reservoir and transmitter of the rabies virus and pose by far the greatest threat to global public health [2,3]. In a true One Health approach [4], elimination of rabies at its animal source would be most impactful and reasonable [5]. To this end, concerted control efforts based on public awareness and mass dog vaccinations were instrumental in rabies control and elimination in Europe and the Americas, where several countries are now considered free from dog-mediated rabies [6]. However, dog-mediated rabies continues unabated in low-resourced countries in Africa and Asia. For various reasons, it represents a challenge to create and maintain adequate herd immunity in dog populations [5], particularly in free-roaming dogs. The latter could be approached by oral rabies vaccination [7].

The Tripartite (FAO, WOAH, and WHO) considers rabies control a priority and in the frame of a global strategic plan, a coordinated, country-centric strategy is followed to eliminate human deaths from dog-mediated rabies by 2030 [8]. This involves national plans for the control of rabies. One of the main hurdles when planning a long-term strategy is the number of dogs to be targeted by vaccination campaigns. Various laborious methods for estimating local dog populations have been described including dog census, distance based methods (transect line counting), foot-patrol transect survey, mark-re-sighting methods, Bayesian models, unmanned Aerial Vehicles (UAV) method, door to door surveys, telephone surveys and KAP studies [9–12]. Because dog population studies are often lacking in countries, estimates are based on a human:dog ratio which is often differentiated between urban and rural populations [13,14]. These calculations are on a country level and can be allocated on the lowest national level of administration. Such administrative units may be quite large in size and eventually heterogenous in their human population density. To overcome these limitations from administrative boundaries, gridded human population datasets have been used, with spatial resolutions constantly improving from several km, to grid cells with a length of 100m [15]. More recently, a method based on machine learning was used to create population maps from satellite imagery at a global scale, with a spatial sensitivity corresponding to individual buildings [16]. This so called High Resolution Settlement Layer (HRSL) has a resolution of 30m and assigns human population to individual buildings. It was validated and shown to be very successful in identifying building footprints [17].

Here, we aim at an optimization of future dog mass vaccination campaigns in the Northern Communal Areas (NCAs) of Namibia by demonstrating the use of HRSL in the context of dog population estimation and analyses of the effectiveness of the mass dog vaccination campaigns using the Oshana region as an example. In the latter, we were particularly interested in whether the vaccination coverage of the dog population could be improved if static vaccination points were strategically set up differently compared to the traditional ones. Additionally, we calculated and compared resulting costs of the different static point vaccination strategies.

## 2. Materials and Methods

### 2.1. Study area

Oshana (18.4305° S, 15.6882° E) is one of the eight regions in the NCAs comprising 8647 km^2^ and has a human population of 175,000 (20 inhabitants/km^2^). The towns of Oshakati, Ongwediva and Ondangwa, all located in the northern part of this region, form a cohesive urban cluster and a centre of important economic activities representing the second largest population concentration in the country after the capital Windhoek. In this area, the implementation of the national dog rabies elimination program was piloted in March 2016– March 2017 [18].

### 2.2. Data sources

HRSL data was downloaded as raster files (.tiff) from the Humanitarian Data Exchange (HDX) platform (https://data.humdata.org/) for Namibia and Malawi, respectively. Additional data on administrative boundaries, schools and cattle vaccination points (crush pens) for the NCAs in Namibia were retrieved from various sources (S1 Table)

### 2.3. Software and GIS analyses

For geospatial analyses both the open-source software QGIS (v.3.16, QGIS.org) and the commercial software ArcGIS ArcMap 10.8.1 (ESRI, Redlands, CA, USA) were used. Initially, the raster file was converted into a vector map, i.e. each identified building was given point coordinates (WGS 84) with the human population associated with this point. This data was transferred into a CSV file for further analyses. Cumulative data were calculated using EXCEL (Microsoft Corporation, Redmond, WA, USA) spreadsheets and visualized in GraphPad Prism v9.0 (GraphPad Prism Software Inc., San Diego, USA).

### 2.4. Estimation of the dog population

A column “dog_pop” was added to the CSV file from the Oshana region. The value for “dog_pop” was calculated using the following parameters: grids cells (30×30 m) with a human population of 0-5 people/grid were assigned to the class “rural remote” with the human population multiplied with a factor of 0.25; i.e a human:dog ratio of 4; “rural dense” (>5-10 people/grid): 0.15; human:dog ratio of 6.6; “semi-urban” (>10-15 people/grid) and “urban” (>15 people/grid): 0.1, human:dog ratio of 10. The ratio of humans to dogs in rural and urban areas used for the calculations of dog population in each grid cell was derived from a recent KAP study and publications from comparable African settings [19,20]. For validation of the approach we used dog census data from Blantyre, Malawi and Namibia [21,22].

### 2.5. Comparison of different static point vaccination strategies

To evaluate the effect of strategic choice of static vaccination points (SVP) on the effectiveness of mass dog vaccination campaigns, we compared the use of traditional crush pens (cattle vaccination sites), pilot study vaccination sites, schools and a combination of schools and pilot study vaccination sites in terms of vaccination coverage of the dog population. Generally, we assumed in our estimates that the compliance to bring dogs to a central vaccination point decreases with distance, as shown before [23,24]. For analysing the vaccination coverage of the local dog population, we buffered the vaccination points with distances (Tab 1, S2 Table) and derived the respective dog population per distance by counting the dogs per grid cells in the areas (see above), and multiplied by the respective assumed compliance of 90-5% (Table 1).

**Table 1:** The assumed relative additive compliance to bring dogs to a central vaccination point.

| distance from vaccination point | compliance to vaccinate dogs |
| --- | --- |
| 0-1 km | 90% |
| 1-2.5 km | 60% |
| 2.5-5 km | 30% |
| 5-10 km | 5% |

As for the comparative cost-effectiveness analysis of the individual strategic approaches, we used vaccination logistic parameters and associated costs, e.g. costs per team/day, number of vaccinations/team/day, vaccination per person/day, vaccine costs as well as distance between points (km), costs per km, cost speed (km/h), staff salary/day, and time for vaccination (in min), from the national dog rabies elimination program and an ORV field trial in the Zambesi region (S2 Table) [25].

## 3. Results

### 3.1. Estimating the population density of dogs

Using different parameters, the population of dogs was derived from the spatially resolved human data sets (Fig 1, 2). The majority of people in Oshana (72.3%) lives in areas classified as rural dense, and less than 10% live in urban or semi-urban areas. As for the dog population, here, most dogs (79.6%) live in rural remote areas (Fig 3). The estimated total number of dogs for the Oshana region ranged between 37,007 when a general human:dog ratio of 5.0 was assumed, and 40,571 when the grid-based gradient was used, and 63,171 if one dog was assigned for each grid cell.

**Fig 1.**
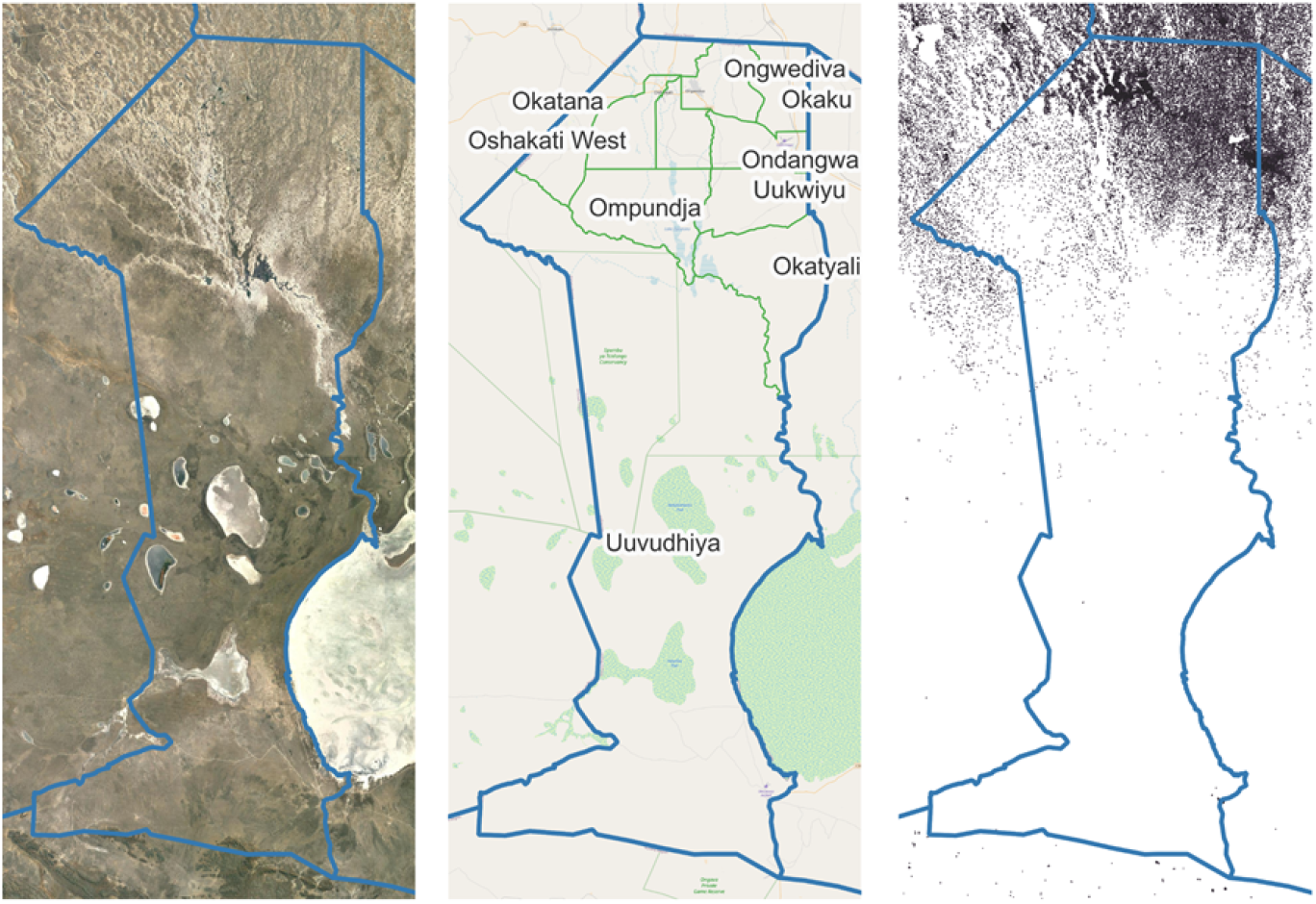
Outline of the Oshana region in Namibia (blue) and details as available from Google earth © (left), the respective administrative boundaries of the constituencies in the Oshana region, with Open Street Map (OSM, OpenStreetMap and OpenStreetMap Foundation; https://www.openstreetmap.org) data as background (middle), and the display of identified buildings/households from the High Resolution Settlement Layer (HRSL), depicted as points.

**Fig 2.**
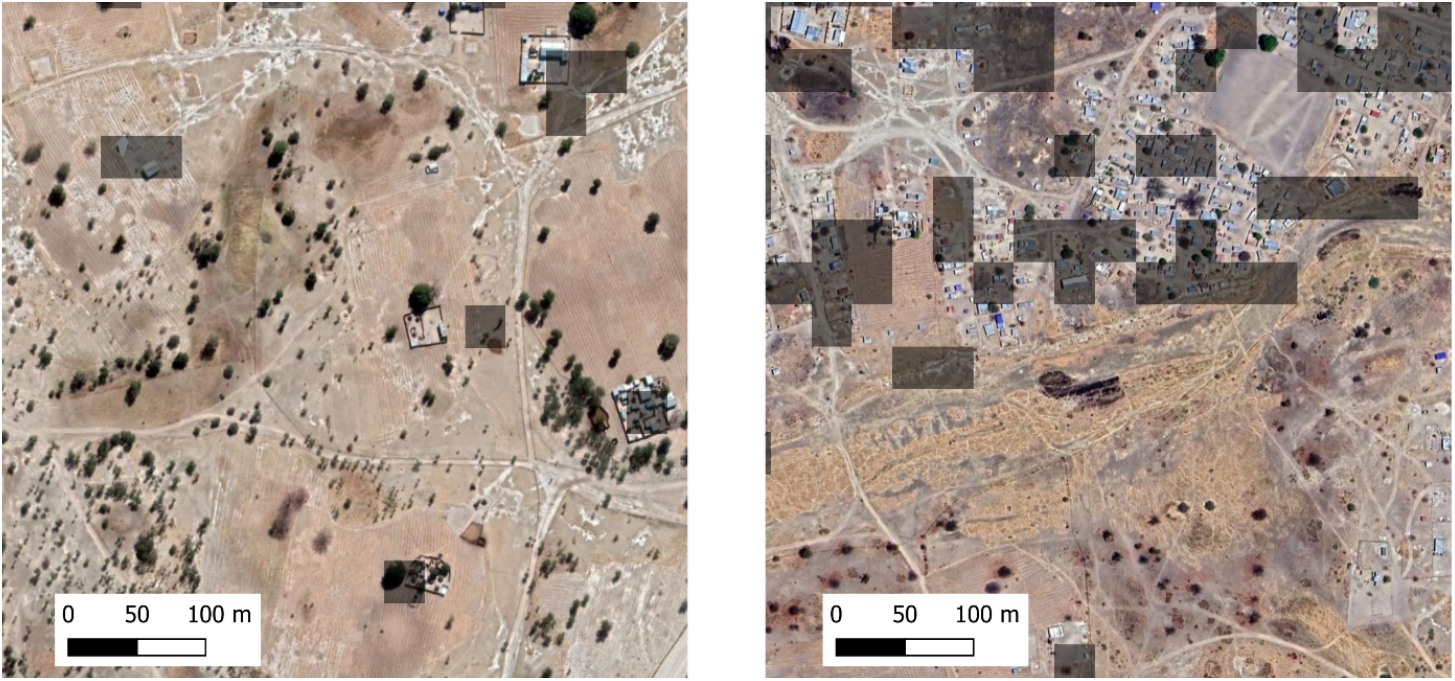
Examples of identified buildings/houses (grey grid cells – 30×30 m) from the High-Resolution Settlement Layer (HRSL) for a rural area in Oshana (left) and a peri-urban area (right). The underlying satellite image of the two depicted sample areas is based on data from Google Earth ©. While the degree of spatial correspondence between the grid cells and the buildings on the satellite images is almost perfect in rural areas, the correspondence in urban areas may be limited but still sufficient.

**Fig 3:**
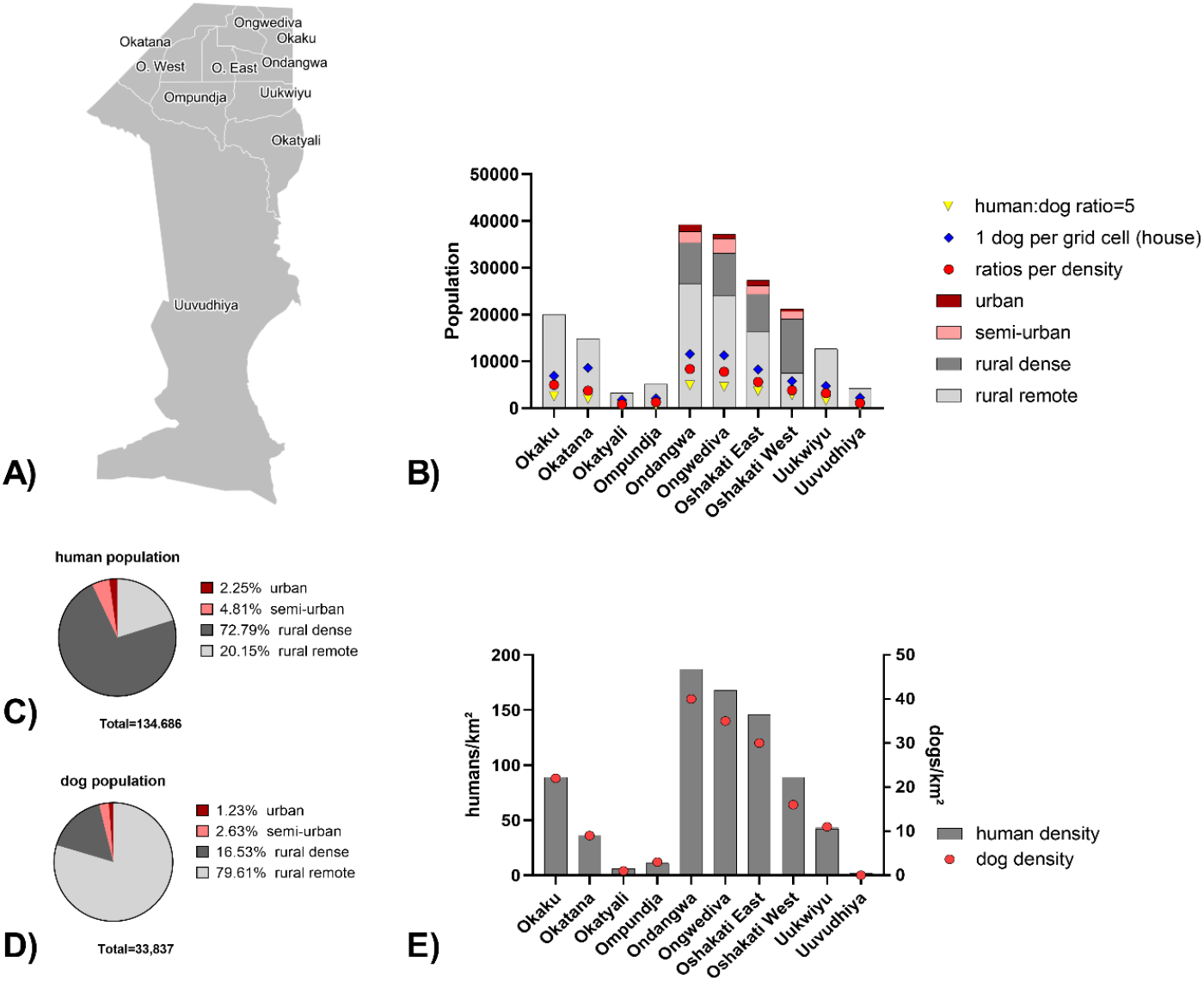
Display of constituencies in Oshana (A) and the estimated number of dogs per constituency depicted based on a fixed human:dog ratio of 5, one dog per identified house, and as per ratios per density class as per HRSL data (B). Proportion of settlement types (based on human density per identified house) of the human (C) and thereof derived dog population (D) in the Oshana region. Human and dog population densities per individual constituency level (E).

The average dog density per km^2^ in constituencies ranged from near zero (Uuvudhiya) to 40 (Ondangwa, Figure 3E). In detail, mapping of dog densities in Oshana showed large areas with high numbers of dogs per grid cells (2.86-4.82) and a rather heterogenous dog density per km^2^, with highest densities close to the major urban areas (Figure 4).

**Figure 4:**
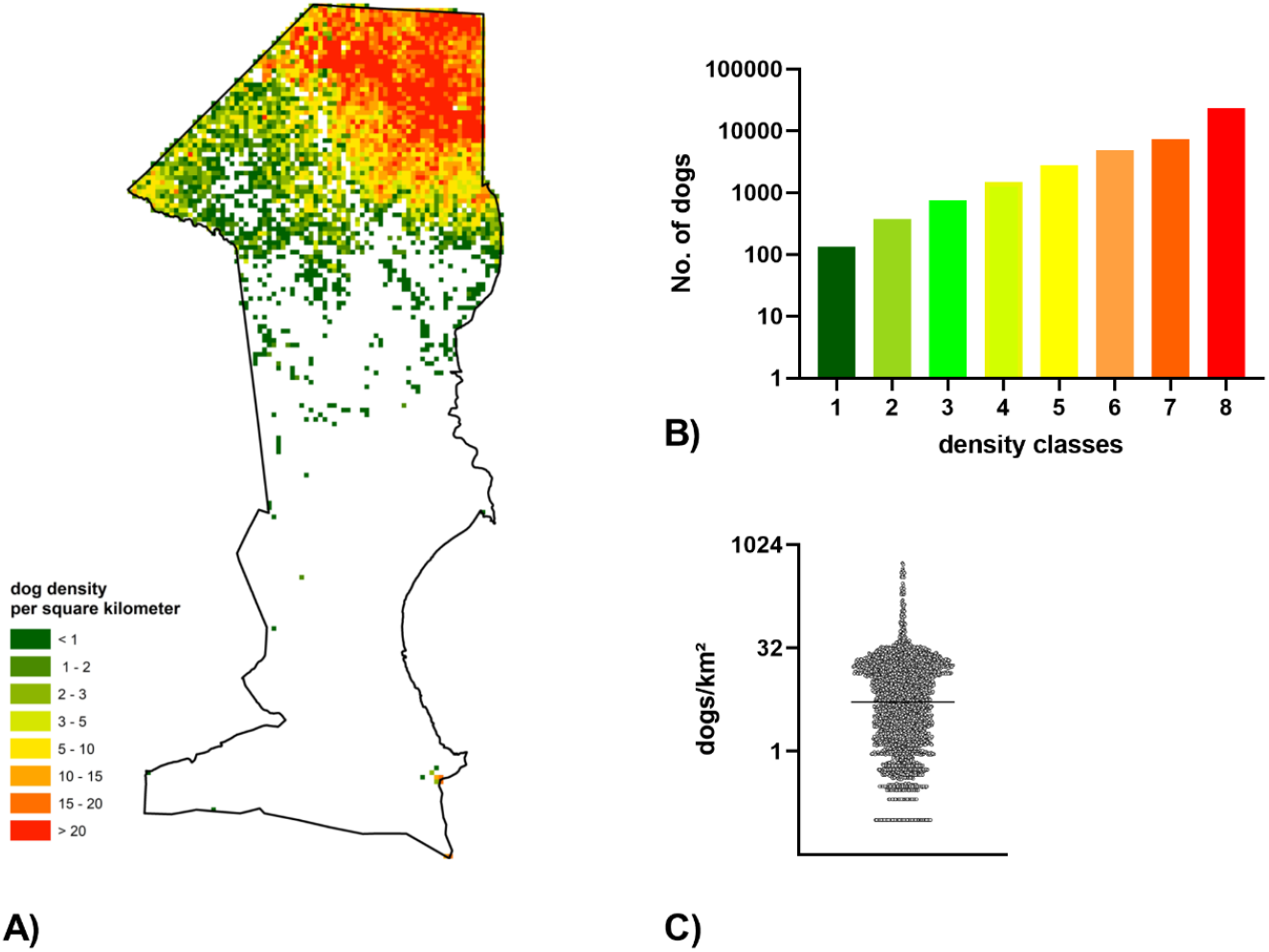
Display of the dog density per km^2^ in Oshana (A). The total number of dogs per class (B), and the overall distribution of dog densities in Oshana (C) is shown.

#### Validation of the approach

We used Blantyre, Malawi, as an example where already good estimates for the dog population existed, based on previous surveys and post vaccination monitoring [21] to validate our approach. When the dog population was based on a literature human:dog ratio of 1:20, the number of dogs was estimated as 38,897, but was higher (46,065) when extrapolated from vaccination and sightings [21]. With remote extrapolation and a slight modification of the parameters, the estimate was 48,017 dogs, only 4% deviation from the actual count. The resulting dog densities show a rather heterogenous pattern, particularly in Blantyre city.

Similarly, the dog population was assessed for four reference areas in the Oshana region and compared with previous estimates from a recent study in Namibia, the applicability of this approach was confirmed [22] (S1 Fig).

Using the gradient for estimation of dogs, and different dog rabies vaccination strategies the potential vaccination coverage of the local dog population was assessed. The vaccination coverage ranged between 44% for vaccinations at cattle crush pens 65% for a combination of schools and cattle crush pens (Figure 5, Table 2).

**Table 2:** Results of different static point vaccination strategies, with their estimated vaccination coverage and costs. The costs are either based on team days (Calculation A) or on individual times and costs for static vaccination point, transport, etc. (Calculation B). The details can be seen and modified in S2 Table.

| Placement of static rabies vaccination points | No. of SVP* | No. of vaccinated dogs | Coverage | Calculation A |  |  | Calculation B |  |
| --- | --- | --- | --- | --- | --- | --- | --- | --- |
|  |  |  |  | Team days | Campaign | Costs per dog | Campaign | Costs per dog |
| Schools | 99 | 24798 | 61% | 496 | \$156.229 | \$6,30 | \$166.213,16 | \$6,70 |
| Crush pens | 54 | 17771 | 44% | 355 | \$111.955 | | \$117.577,58 | \$6,62 |
| Combination Schools/Crush pens | 153 | 26437 | 65% | 529 | \$166.556 | | \$181.490,84 | \$6,86 |
| Vaccination Sites + Health facilities | 52 | 20298 | 50% | 406 | \$127.875 | | \$133.422,69 | \$6,57 |

\*SVP: static vaccination points

**Figure 5:**
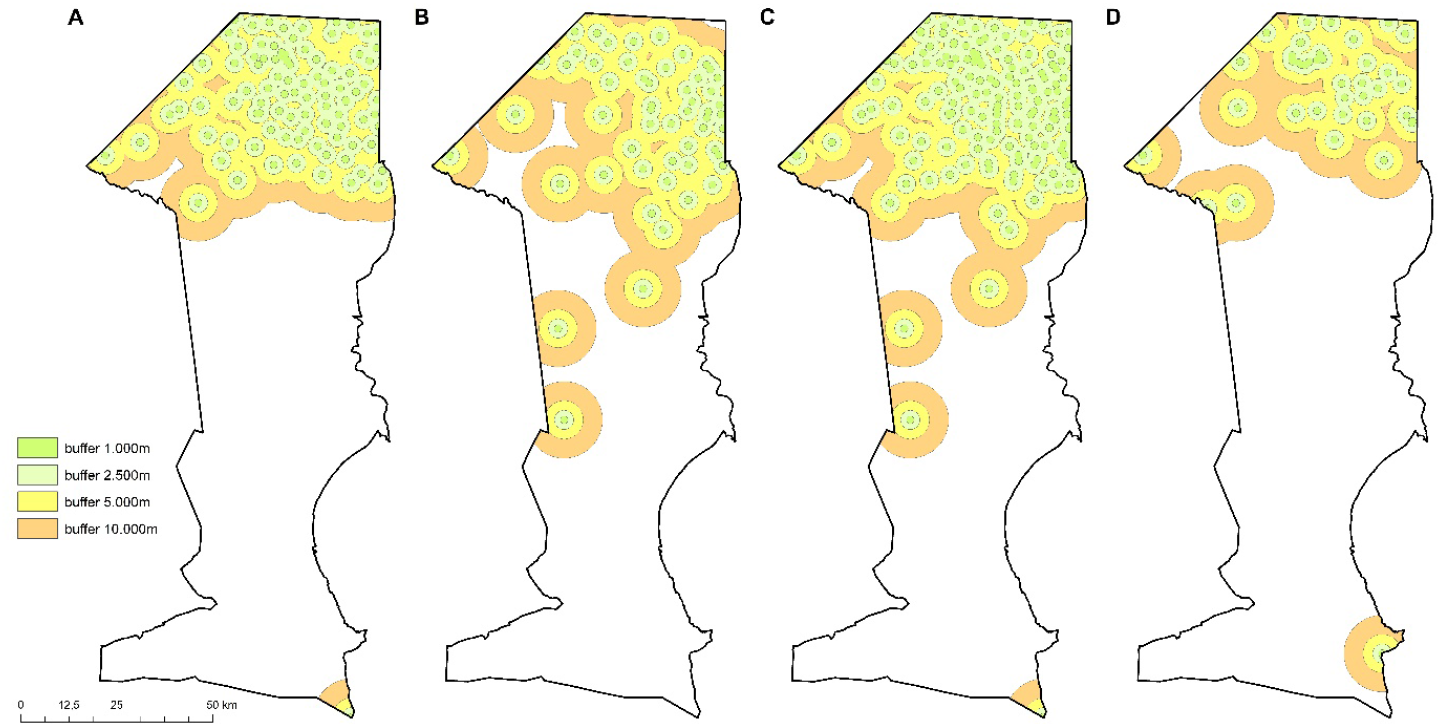
Comparison of different static point vaccination strategies: Display of the estimated spatial vaccination coverage per strategic approach, (A), schools (B), crush pens (cattle vaccination posts, C) a combination of schools and crush pens (cattle vaccination posts) and D) human vaccination sites and health facilities

In terms of cost-effectiveness of the different static point vaccination strategies, the resulting costs per dog vaccinated ranged between US$6.30 when a uniform rate per team and day was assumed and US$6.86 when time for travel and individual vaccination was separately calculated (Tab 2; S2 Table). In this approach, vaccination close to schools would be most cost effective. In both calculations, vaccinations at crush pens had the lowest overall costs per campaign.

## 4. Discussion

Detailed and up-to-date spatial datasets that accurately describe the distribution of human populations can be used for measuring the impact of population growth, monitoring changes, environmental and health applications, and planning interventions [26]. While Facebook data for good have been used for analyzing disaster management [17] there benefit for epidemiological studies and veterinary public health intervention strategies has not been explored yet.

As shown in this study, the availability of high-resolution human population data at a scale of 30×30m can even contribute to better spatial analyses of local dog populations, and thus support global efforts in fighting dog-mediated rabies e.g. optimization of dog rabies vaccination campaign planning and monitoring. In this example from the Oshana region, we demonstrated the utility of estimating the dog population (Fig 3). Inevitably, there has to be a rough indication of the human:dog ratio to infer the dog population, and here we fortunately had already available estimates from a previous study [22]. However, in the absence of such data, published data from neighboring countries or similar socio-economic settings could initially be used as a proxy [13,20], until further data is available.

The average predicted dog density for the entire Oshana region was 8.2 dogs/km^2^ which is similar to the 8.8 dogs/km^2^ estimated for Tanzania [27]. This density in Oshana was clustered and varied greatly between constituencies (Fig 3) and between individual 1km grids, with densities ranging between 0 and 551 dogs/km^2^ (Fig 4). Although the dog density in Oshana on average is much lower than dog populations in densely populated areas in South-East Asia, e.g. Thailand (251.6 heads/km^2^), Philippines (468 heads/km^2^) [28,29], surveillance data demonstrate that this density is still sufficient to maintain a rabies transmission chain among dogs [30], which make concepts to concentrate control efforts to more densely populated areas a threat to a real elimination strategy [31].

Besides this similarity in dog densities in Tanzania and Oshana, using another example of Blantyre we could show, that this approach is applicable to other settings as well, with slight modifications to the parameters. Of course, the more data is available, e.g. on differences in dog ownership in different regions, the better is the population estimate. Thailand used surveys and developed a model to estimate their dog population [28]. While this approach may well be more precise, it needed a lot of effort and GIS expertise, which may not be available in most rabies endemic countries.

In this example, we have used a gradient for the population estimate based on human density per grid cell, with a higher human density (as a proxy for urbanization) having higher human:dog ratios. The estimates of a simple generalized human:dog ratio did not differ much from our method and it is likely that this can be more easily applied for program managers.

Also, it has to be mentioned that even with surveys, e.g. a household level or at schools, a rather variable number of dogs is estimated, that only partly matches with sight-resight studies [27]. Such dog population studies on the ground are complex and costly, and using our remote-sensing approach, rabies program managers are better capable of estimating dog populations, plan vaccination campaigns and use the number of dogs vaccinated in a given area to easily extrapolate the vaccination coverage as demonstrated before [32,33], without the need for post-vaccination surveys.

The application for spatial analyses of the effectiveness of static point vaccination strategies in the Oshana region in Namibia demonstrated that standard approaches (i.e. in Namibia, vaccination of dogs at crush-pens) have inherent limitations in their outreach to the people, particularly in this dispersed human settlement (Tab 2). This finding confirms results from an earlier study, during which gridded population data in four regions of the NCAs including Oshana revealed a suboptimal vaccination coverage in the great majority of grid cells (82%) with a vaccination coverage below 50% during the vaccination campaigns 2019 and 2020 [32]. Based on our analyses, vaccination at human health infrastructures and near schools would increase the vaccination coverage, but only switching to a combination of vaccination points at crush-pens and near schools would bring the coverage closer to the 70% threshold at which there is a high likelihood that the transmission chain is interrupted [30,34]. However, these marginally increased percentages are based on ca. 50% more vaccination points associated with higher efforts.

In the absence of sufficient funds to cover all areas in this way, the high spatial resolution of dog densities, among other factors, may guide campaign managers to strategically apply the resources they have, as recommended before [35].

Our calculations on vaccination coverage are based on fairly optimistic assumptions, e.g. that most dogs within a 1km radius to a vaccination point will be vaccinated. Data from Malawi suggests that the compliance even close to the vaccination point is close to the 70% and rapidly decreases [24]. If these parameters are used, the vaccination coverage decreases further and is closer to the real-life estimates from the recent Namibian dog vaccination campaigns that failed to reach the goal of 70% [32]. With the high percentage of free roaming dogs in the Namibian NCAs the challenge to reach enough dogs with central-point vaccinations could already be demonstrated using CDC’s vax calculator [36]. With the data from the Oshana region entered, the maximum coverage reached is 66%. This exemplifies that tools and data are available that can help to optimize rabies vaccination programmes. While the vax calculator makes general cost and vaccine coverage calculations, the use of HRSL can be applied in the context of spatially resolved dog population estimation and subsequent planning and analyses.

Even if distances to vaccination points were lowered, people were aware of the vaccination campaign and would be willing to bring their dogs, handling of dogs and the availability of the owners would still reduce the compliance to bring dogs to vaccinations, demonstrating the inherent limitations of static point vaccinations [7]. Door-to-door vaccinations are no real alternative, as the spread-out human population make this approach too costly. In general, because of the logistical constraints of long distance travel, the costs to vaccinate a single dog is higher than in most other settings elsewhere [36]. In our setting the costs was USD 6.3, which is similar to cost estimates from Tanzania [37]. Given the limitations of static point vaccinations and the advantages in the effectiveness of ORV [7,38], particularly in free roaming dogs [39], a shift to ORV would not only increase the vaccination coverage particularly in those dogs that are hard-to-reach for parenteral vaccination, but could also save sparse resources [25].

## 5. Conclusions

Freely available high-resolution population datasets can be an invaluable source of information, particularly in rapidly changing environments. This information can be used to estimate the dog population on a spatial resolution that allows for fine-scale planning and evaluation, thus saving resources that would otherwise be directed in e.g. post-vaccination monitoring as recommended before [27]. These datasets can be visualized and analyzed even using open-source GIS software packages, e.g. QGIS, and therefore are a promising tool, particularly for resource-limited settings. Also, the high resolution of settlement structures and human populations can be used for disease modelling purposes [40] and will perfectly integrate with other GIS data gathered during vaccination programs [41,42] and surveys [21,24]. The wide utility of the HRSL was demonstrated for rabies here, but it is only an example and can be used form many more applications, particularly in the field of public health.

## Data availability

All data is either part of this publication, publicly available or published as supplementary file.

## Acknowledgments

This study received no external funding. Generally, Namibia’s animal disease control efforts, including rabies, are gratefully supported by funds from the German Federal Ministry of Food and Agriculture and the German Federal Ministry of Economic Cooperation and Development through projects coordinated by the World Organization for Animal Health (WOAH). We also thank Daike Lehnau (FLI) for GIS-assistance and Nicolai Denzin (FLI), Ronald Schröder (FLI) and Fred Lohr (Mission Rabies) for constructive discussion.

## Supporting information

S1 Table: Excel-Spreadsheet for the calculation of vaccination coverage and costs.

S1 Figure:

**S1 Figure:**
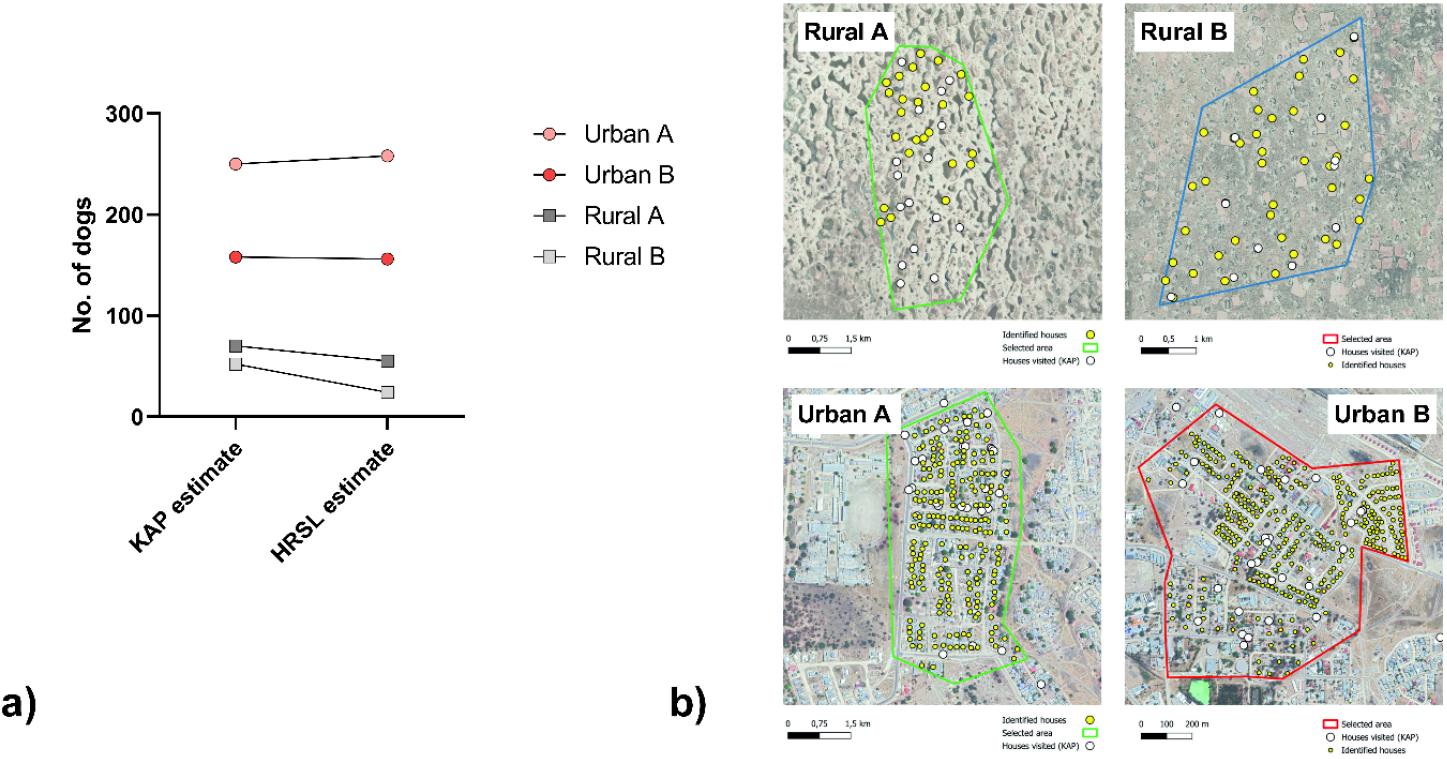
a) Comparison between dog population estimation based on a recent KAP study and the respective estimate using High-Resolution Settlement Layer (HRLS). The maps of the reference zones used for validation are shown (b).

